# Computational pipeline for the PGV-001 neoantigen vaccine trial

**DOI:** 10.1101/174516

**Authors:** Alex Rubinsteyn, Julia Kodysh, Isaac Hodes, Sebastien Mondet, Bulent Arman Aksoy, John P Finnigan, Nina Bhardwaj, Jeffrey Hammerbacher

**Affiliations:** Department of Genetics and Genomic Sciences, Icahn School of Medicine at Mount Sinai, New York, NY 10029; Tisch Cancer Institute, Icahn School of Medicine at Mount Sinai, New York, NY 10029; Department of Microbiology and Immunology, Medical University of South Carolina, Charleston, SC 29425

**Keywords:** neoantigens, personalized vaccine, immunoinformatics, genomics, computational pipeline

## Abstract

This paper describes the sequencing protocol and computational pipeline for the PGV-001 personalized vaccine trial. PGV-001 is a therapeutic peptide vaccine targeting neoantigens identified from patient tumor samples. Peptides are selected by a computational pipeline which identifies mutations from tumor/normal exome sequencing and ranks mutant sequences by a combination of predicted Class I MHC affinity and abundance estimated from tumor RNA. The PGV pipeline is modular and consists of many independently usable tools and software libraries. We draw attention to three particular tools which may be useful to other groups working on neoantigen vaccination. (1) Epidisco is a workflow which orchestrates the parallel execution of the PGV pipeline, including both common steps such as alignment as well as tools which have been developed specifically for the PGV-001 trial. (2) Vaxrank uses somatic variants and tumor RNA reads to select peptides for inclusion in a patient’s vaccine. (3) Isovar is a library used by Vaxrank to determine the mutated protein sequence associated with a genomic variant from supporting tumor RNA reads. We hope that the functionality of these tools may extend beyond the specifics of the PGV-001 trial and enable other research groups in their own neoantigen investigations.

## Introduction

Cancer neoantigens are antigens presented on tumor cells which, due to either mutation or modification, are not presented on normal cells. Neoantigens generated by tumor DNA mutations have been shown to play a significant role in mediating the destruction of tumor cells by the adaptive immune system ^1–3^. Several groups have used therapeutic vaccines targeting neoantigens to clear tumors in murine models ^4–6^. Consequently, many human neoantigen vaccine trials are now under way and several have even published promising early results^7,8^. Since very few cancer mutations are recurrent between patients, the identification of neoantigens requires a personalized genomic approach ^9^. We describe the sequencing protocol and immunogenomic pipeline of PGV-001, a neoantigen vaccine trial at the Mount Sinai Hospital^10^.

The PGV (Personalized Genomic Vaccine) pipeline takes tumor/normal sequencing data as an input and generates a ranked list of mutated peptide sequences. The various steps along the way of determining a personalized vaccine’s contents are implemented as configurable independent tools. We highlight several of these tools (Isovar, Vaxrank, and Epidisco) since they may be potentially useful to other groups working on similar neoantigen vaccine trials.

## Overview of the PGV-001 Personalized Vaccine Trial

PGV-001 is a phase I clinical trial at Mount Sinai Hospital, studying the safety and immunogenicity of a multi-peptide personalized genomic vaccine for treatment of cancers. A PGV dose consists of 10 synthetic long peptides ^11^, each containing a somatic mutation from the patient’s tumor, as well as an immunostimulatory adjuvant: Poly-ICLC ^12^. In the PGV-001 trial, the personalized vaccine is administered in the adjuvant setting, for patients who undergo a complete resection and have no evidence of residual disease. The personalized vaccine is administered as an intramuscular injection and is given to the patient 10 times over a span of 6 months.

The inputs to the PGV pipeline (see Figure 2) are unmapped sequencing data from tumor DNA, tumor RNA, and “normal” patient DNA, typically from peripheral blood. The two DNA samples are compared to identify somatic mutations which are unique to the tumor. The RNA sample is used to quantify the degree to which a mutation is expressed and to determine the mutated protein sequence.

**Figure 1:**
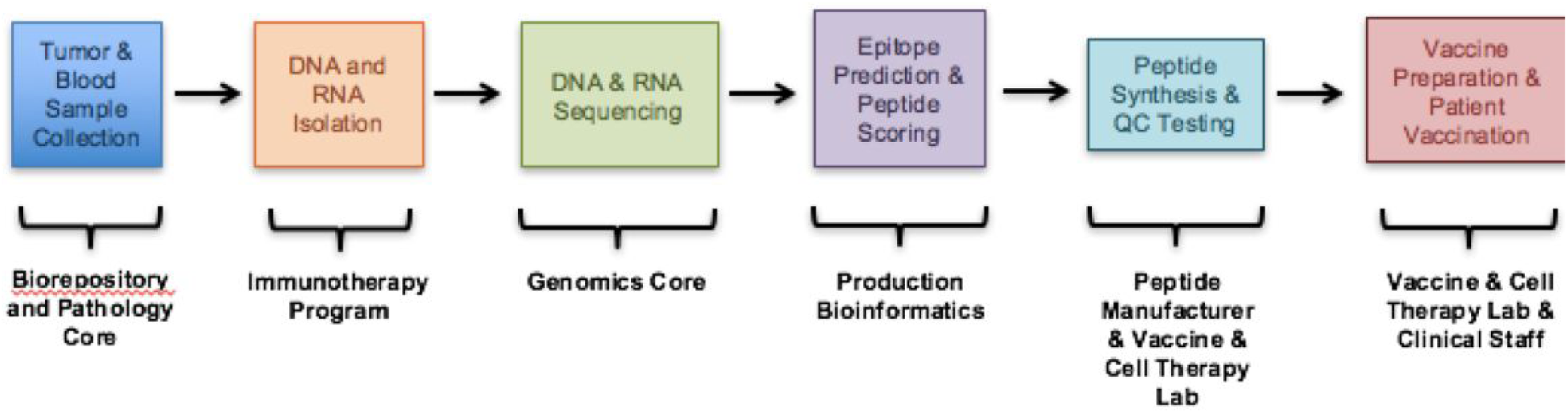
Overview of PGV-001 Trial.

**Figure 2:**
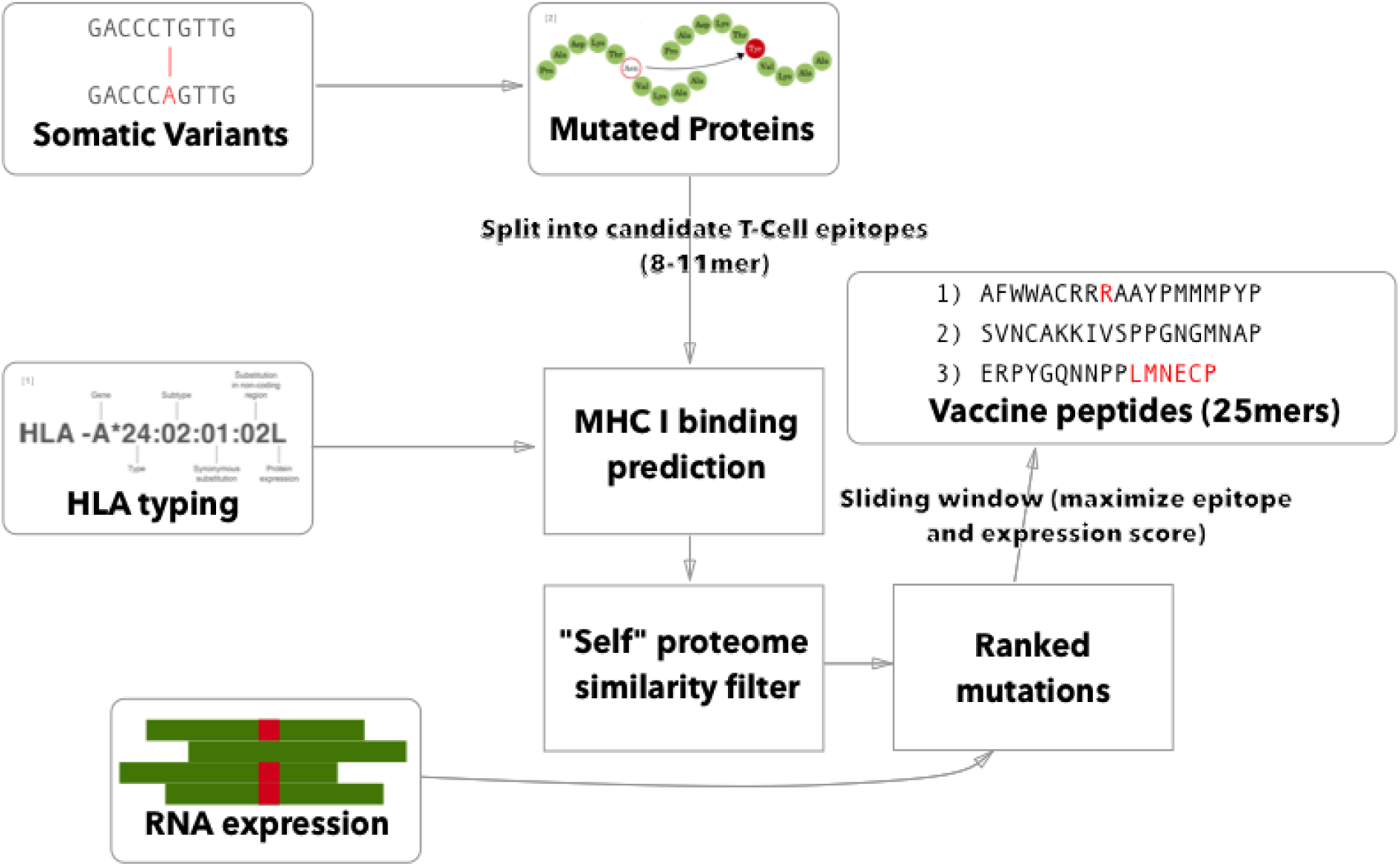
Overview of PGV Computational Pipeline.

To make immunological predictions from these mutated protein sequences, it is necessary to know the patient’s HLA type. This can be determined computationally from exome or bulk RNA sequencing or validated externally using HLA-specific amplicon sequencing ^13^. The PGV pipeline currently uses seq2hla^14^ for HLA typing from tumor RNA while also using amplicon sequencing of normal DNA for validation. Thus far, across 10 patients, the two methods have only disagreed on a single allele. This high degree of concordance matches our previous experience with HLA typing of fresh frozen tissue samples; formalin fixed tissue is more likely to give discordant results between different sequencing methods.

After the patient’s HLA type is determined, we use NetMHCpan to predict the binding affinity of mutated peptides to all patient class I alleles. It should be noted that the pipeline is written to freely allow the use of other predictors, such as NetMHC^15^ and MHCflurry^16^. We then combine affinity predictions with an estimate of each mutated protein’s abundance to generate a ranked list of vaccine peptides. The manufacturer then attempts to synthesize many different peptides from this list and delivers the top 10 sequences which they are able to purify to sufficient quality and quantity. The lyophilized peptides are dissolved in DMSO and mixed with poly-ICLC immediately before injection.

## Sequencing protocol for DNA and RNA

The sequencing protocols used for both DNA and RNA can dramatically affect the sensitivity of variant detection, and thus ultimately change the results of the vaccine pipeline. The largest determinants of sensitivity are the sample quality, method of sequencing library preparation, and quantity of sequenced reads. Whenever possible, PGV uses fresh frozen tumor tissue samples, which results in significantly improved variant detection accuracy as compared with sequencing of formalin fixed (FFPE) samples^17^. An additional benefit of using fresh frozen samples is that mRNA can be enriched using poly-A capture, whereas the fragmented RNA of FFPE samples can only be prepared with less efficient methods such as ribosomal depletion ^18^. For patients with solid tumors, normal DNA is extracted from peripheral blood rather than potentially contaminated adjacent tissue ^19^.

Fragmentation by sonication was preferred to transposase-based methods^20^ due to significant sequence bias, leading to lost coverage after marking duplicate reads^21^. Among the exome enrichment techniques which use sonication, we chose Agilent’s SureSelect XT kit due to its efficient rate of capturing on-target reads^22^.

We chose to target 150X mean coverage for the normal DNA (exome) sequencing since this was found to be the point of diminishing sensitivity across different variant calling pipelines ^23^. Several of the cancer types allowed in the PGV-001 trial (particularly lung and head/neck cancers) have been shown to result in systematically low purity samples^24^. To accommodate a significant degree of non-cancerous DNA, we assume 50% tumor purity and consequently target 300X exome coverage for the tumor DNA sample.

A final consideration is the choice of read length, which does not significantly impact variant discovery from DNA but does impact variant phasing in RNA. Since a 25mer vaccine peptide is translated from 75bp of coding sequence, PGV could theoretically use any read length above that minimum. To allow for many distinct aligned positions overlapping the same region of coding sequence, the PGV protocol uses 125bp reads. These provide a good compromise between length and base quality on the HiSeq 2500 instrument. Newer sequencing platforms from Illumina allow longer reads (150bp) without significant increases in base error.

## Somatic variant calling

The tumor and normal DNA samples are aligned against the human GRCh37 reference genome using BWA-MEM^25^. The tumor RNA is aligned using STAR^26^, which has been found to have particularly high sensitivity for detecting indel variants^27^. The alignment files were processed according to GATK Best Practices^28^. To improve the sensitivity of indel calling, we also perform assembly-based realignment on RNA reads containing potential indel variants. Somatic variant calling is then performed using Mutect^29^ and Strelka^30^, whose results are combined by taking a union of called variants. In cases where Mutect and Strelka yield an insufficient number of variants, we also use Mutect2 to increase sensitivity. Once all these steps have completed, we have combined VCF of candidate somatic variants and aligned RNA reads which will be used to gauge expression of these variants and determine mutated coding sequences.

## Isovar: determining the mutant protein sequence

There are several different software packages which predict the protein-level effect of a coding mutation^31–33^. However, for the purposes of selecting a vaccine peptide’s sequence, it is not sufficient to predict a DNA mutation’s protein effect in without considering the transcripts in which it occurs. A somatic mutation can be associated with selective splicing of particular RNA isoforms^34^ and can also co-occur with other genomic variants. Thus, the tumor RNA sequencing data is also used to determine a mutant coding sequence.

For each mutation, it is possible to infer multiple coding sequences from supporting RNA reads due to sequencing error, splicing diversity, and tumor heterogeneity. To account for these potentially complicating factors, we developed a tool called Isovar^35^, which can be downloaded from https://github.com/hammerlab/isovar. Isovar uses RNA to assemble the most abundant coding sequence for each mutation. An overview of the algorithm is given in Figure 3.

**Figure 3:**
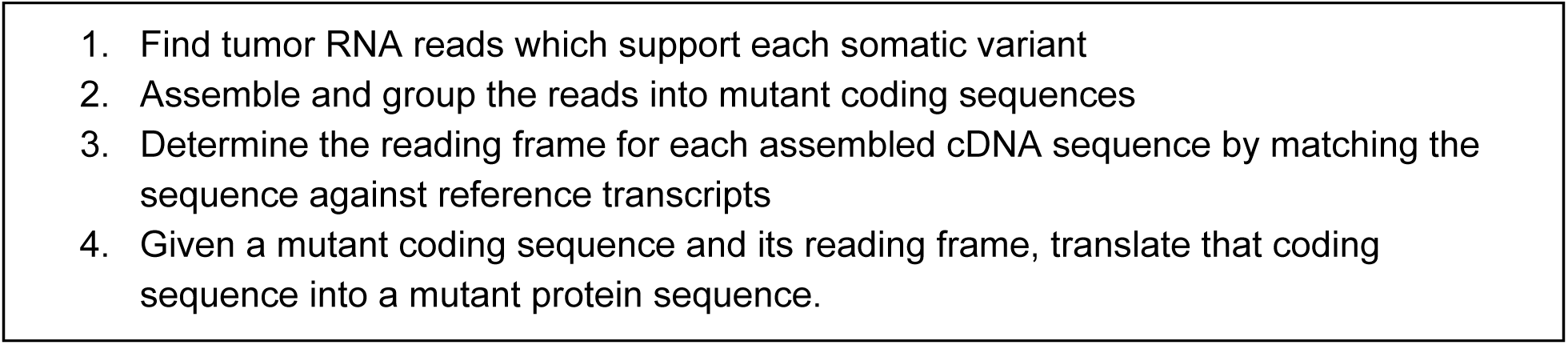
Overview of Isovar Algorithm for Determining Mutant Protein Sequence.

**Figure 4:**
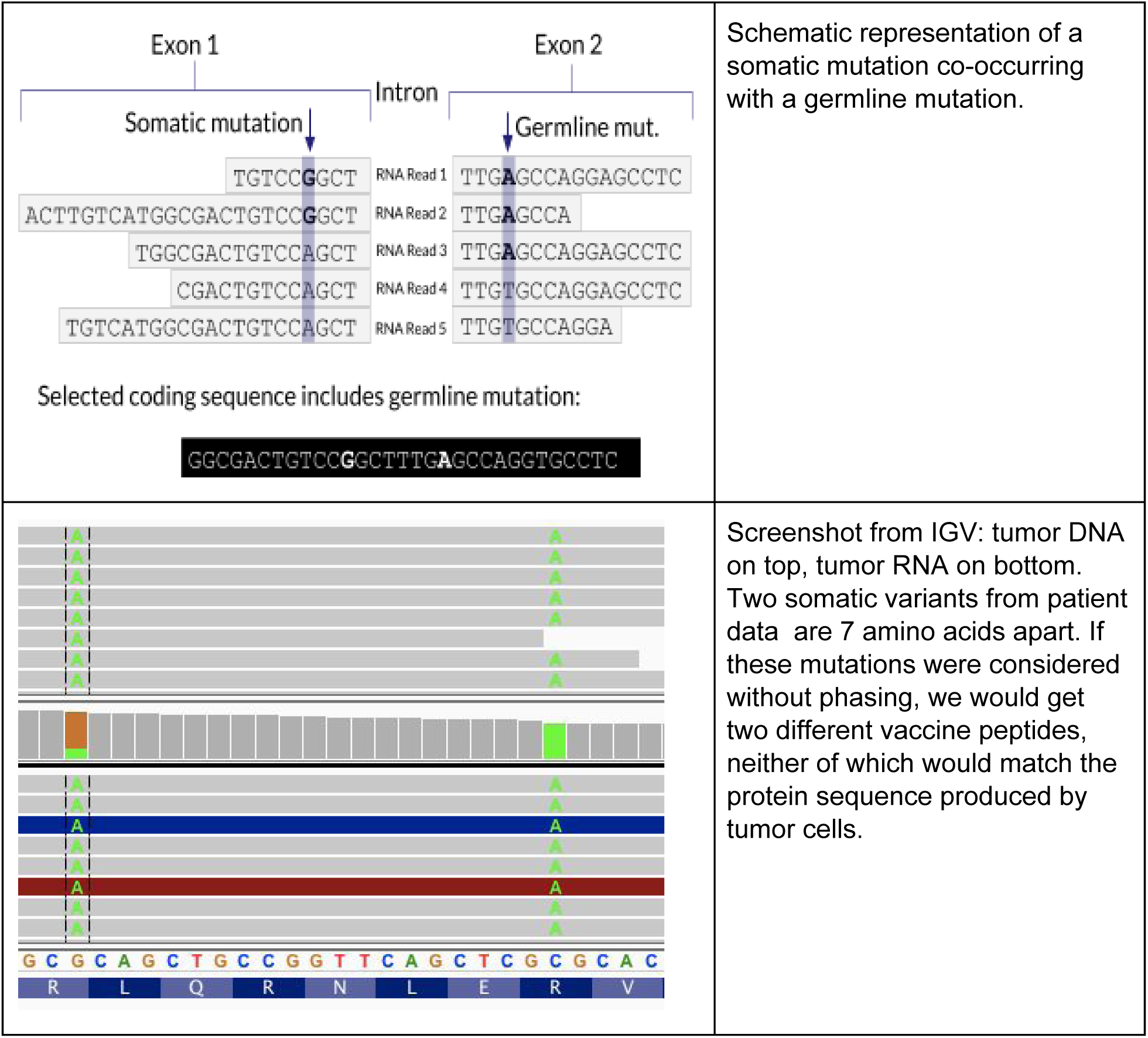
Phasing Adjacent Variants.

One advantage of using RNA to determine the coding sequence is that it phases adjacent (germline or somatic) variants. A further advantage is that Isovar, by using mutation-supporting RNA reads to determine each mutant protein sequence, naturally estimates allele-specific expression. If the PGV pipeline, on the other hand, used bulk expression for each gene it would potentially overestimate how much of a mutant protein is being made. In an extreme case, all of the RNA reads aligning to a particular gene could be wildtype, with none supporting the somatic variant of interest.

## Vaxrank: vaccine peptide selection

Once we have determined the amino acid sequences containing somatic mutations and estimated their abundance in the tumor, the final step is to rank them according to desirability of inclusion in personalized vaccine.

There are many potential correlates of immunogenicity which can be used to prioritize neoantigens, such as expression, MHC binding affinity, peptide-MHC complex stability, proteasomal cleavage, and other antigen processing steps. Of those, the PGV pipeline optimizes for high expression and predicted strong Class I MHC binding. There are several published computational predictors of Class I MHC binding affinity which have demonstrated high accuracy^15,36,37^. PGV uses NetMHCpan^36^ due to its extensive coverage of patient alleles.

The final ranking of candidate vaccine peptides according to predicted MHC binding and expression is performed by a tool called Vaxrank^38^. Vaxrank identifies high-affinity mutant MHC ligands within each peptide and combines these predictions into a single MHC binding score. This score is then scaled according to the expression of that mutation in the tumor. The formula for computing these MHC and expression scores is given in Figure 5. The scale and offset for MHC affinity normalization was determined by a logistic fit of affinity versus immunogenicity from the dataset used to determine the classical 500nM affinity threshold^39^. There is no rigorous justification for the multiplicative scoring function, other than the intuition that epitope abundance and MHC affinity are independent prerequisites for immunogenicity.

**Figure 5:**
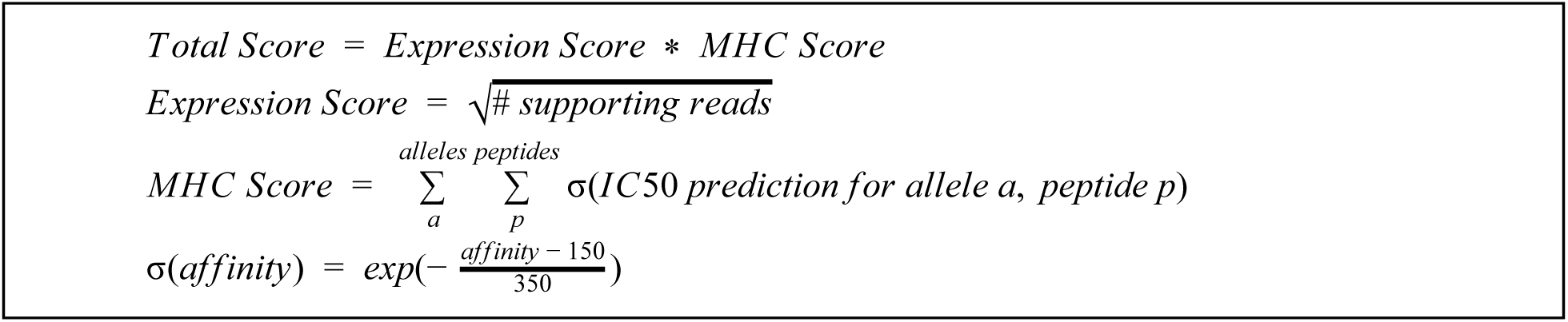
Scoring Criterion for Selecting Vaccine Peptides.

Since some peptides cannot be manufactured using solid-phase synthesis, our vaccine peptide ranking algorithm includes manufacturability heuristics, such as minimization of cysteines. Vaxrank can be downloaded from https://github.com/hammerlab/vaxrank.

## Epidisco: parallel workflow for the PGV pipeline

The actual execution of the PGV computational pipeline requires careful orchestration and combination of various computational steps. Furthermore, the time and the computational power needed to complete each of these intermediate steps within the PGV pipeline varies: for example, somatic mutational calling on the whole genome can easily become a major bottleneck for a pipeline if the details of the execution is not well planned.

We have developed a suite of tools computational tools that enable the PGV pipeline to follow the best practices defined by the community, run faster, and tolerate failures without sacrificing any efficiency. The individual infrastructure tools used by the PGV pipeline are enumerated in Supplementary Table 3. The unifying workflow which implements the logic of PGV is called Epidisco. Epidisco parallelizes a large portion of the somatic mutation calling pipeline by splitting sequencing data by chromosome and running computationally intensive parts of the pipeline on each chromosome. This allows efficient and simultaneous use of tens of computation units towards a single goal and therefore reduces the amount of time and resources required for such demanding computational tasks. Furthermore, QC checks for each data type, the processing of the RNA sequencing data, and the joint analyses of the normal-tumor DNA sequencing data are all run independently, again enabling us to divide the computational tasks needed for each to multiple computational units and run them simultaneously.

Epidisco supports local computation, traditional HPC scheduling, and cloud-based resources from well-known providers. On a typical machine, running the complete PGV pipeline for a single patient can take up to 4 days; but making use of 5 or more computers for parallelization reduces the overall running time down to a single day. The parallelization of the PGV pipeline becomes more prominent once more than one patient is being analyzed simultaneously, which allows efficiently saturating the computational resources and allow the pipeline to scale according to the actual workload. This also make it feasible to use PGV pipeline on a cohort level where fast and efficient pipelines are required to be able to analyze data for tens of patients in a fast, efficient, and accurate way.

Epidisco also makes the PGV pipeline tolerant to failures of intermediate steps and allows resuming the pipeline from the point of failure with a simple restart request. By handling such failures in an automated way, carrying out cleaning procedures, and restarting only the tasks that need to be re-run, the workflow makes it easier for researchers to operate such complex computational tasks. Epidisco provides command line and web-based utilities to facilitate starting a new workflow, collecting the results, and troubleshooting specific parts of a pipeline.

## Discussion

The PGV pipeline is a modular, highly configurable, freely available method for selecting the contents of a therapeutic neoantigen vaccine. The PGV pipeline has been used to predict vaccine peptides for several mouse models (LLC, B16 F1/F10), 5 “dry run” patients whose samples were processed according to the PGV protocol but did not participate in the trial, and 5 patients which were being considered for enrollment in the trial. A summary of the results from the “dry run” patients is presented in Supplementary Table 1. Of the patients eligible for enrollment, 1 has been treated so far and another enrolled. The remainder did not enroll due to progression of disease or low quality tumor samples.

Several other groups have released pipelines for neoantigen vaccine prediction, most notably pVAC-seq^40^ and MuPeXI^41^. A deep comparison between neoantigen pipelines likely requires evaluating T-cell response and anti-tumor activity after vaccination, which is well beyond the scope of this paper. There are, however, a few obvious differences between the PGV pipeline and others which have been published.

- **Modularity:** The PGV pipeline has been developed as a collection of flexible standalone tools, rather than a single monolithic script (see Supplementary Tables 2 & 3). These components are brought together in the Epidisco library, which allows users to run highly configurable genomics and immunoinformatics pipelines. For example, the variant calling and epitope prediction components of the PGV pipeline have also been used through Epidisco for retrospective analyses of checkpoint blockade clinical trials^42^.
- **Inputs are FASTQ files:** MuPeXI and pVAC-seq both require the implementation of separate genomics pipeline to infer patient HLA type, call somatic variants, and quantify expression. The PGV pipeline, by contrast, is self-contained in the sense that its inputs are raw FASTQ files and its outputs are vaccine peptide predictions.
- **Dependence on tumor RNA:** The PGV pipeline relies on tumor RNA reads to determine the mutant protein coding sequence. MuPeXI and pVAC-seq, by contrast, only consider expression data after predicting a mutant protein sequence from a variant in isolation. PGV’s approach has potential benefits in capturing altered patterns of splicing and phasing somatic variants with other nearby variants. These potential benefits, however, have yet to be evaluated systematically.
- **Liberal software license:** All of the software components which comprise the PGV pipeline are freely available under the Apache software license. MuPeXI and pVAC-seq, by contrast, use more restrictive licenses, which might prohibit their integration with other bioinformatics software.
- **Optimization of peptide sequence for solid phase synthesis:** PGV appears to be unique among freely available neoantigen pipelines in attempting to choose peptides whose sequence content is more likely to be successfully manufactured. We have found this to be an important step, especially when using longer peptides, due to the significant delays introduced by failed synthesis or purification attempts.

The PGV-001 trial is the first in a series of planned neoantigen vaccine investigations. Several improvements to the PGV pipeline are planned, including the use of genomic fusions and other structural variants as neoantigen sources, clonality as a consideration for variant prioritization, and additional immunological predictions such as proteasomal cleavage and Class II MHC binding. As immune response data from ongoing preclinical work and PGV-001 becomes available, our method for combining correlates of immunogenicity into a single ranking will likely undergo significant tuning.

---

**Supplementary Table 1:** Summary of Results for Dry Run Patient Samples

|  | Dry Run Patient #1 | Dry Run Patient #2 | Dry Run Patient #3 | Dry Run Patient #4 | Dry Run Patient #5 |
| --- | --- | --- | --- | --- | --- |
| Variants | 501 | 888 | 591 | 663 | 912 |
| Coding | 180 | 253 | 173 | 231 | 305 |
| Predicted MHC ligands | 11 | 9 | 17 | 32 | 22 |

**Supplementary Table 2:** Computational Tools for Genomics and Immunology

| Name | Description | URL |
| --- | --- | --- |
| PyEnsembl | Provides reference genome sequences annotations | github.com/hammerlab/pyensembl |
| Varcode | Predicts coding effects from DNA mutations | github.com/hammerlab/varcode |
| Isovar | Determines mutant protein sequence from called somatic variants and tumor RNA | github.com/hammerlab/isovar |
| Vaxrank | Overall vaccine selection tool with ranking logic | github.com/hammerlab/vaxrank |
| MHCtools | Common interface to many MHC predictor software packages | github.com/hammerlab/mhctools |

**Supplementary Table 3:** Computational Tools for Workflow Infrastructure

| Name | Description | URL |
| --- | --- | --- |
| Ketrew | Resumable workflows | https://github.com/hammerlab/ketrew |
| Biokepi | Wraps bioinformatics tools | https://github.com/hammerlab/biokepi |
| Secotrec | Cluster/node management | https://github.com/hammerlab/secotrec |
| Epidisco | The actual implementation of the PGV pipeline | https://github.com/hammerlab/epidisco |

